# A high soluble-fibre allele in wheat encodes a defective cell wall peroxidase responsible for dimerization of ferulate moieties on arabinoxylan

**DOI:** 10.1101/2023.03.08.531735

**Authors:** Rowan A.C. Mitchell, Maria Oszvald, Till K. Pellny, Jackie Freeman, Kirstie Halsey, Caroline A. Sparks, Alison Huttly, Sebastien Specel, Michelle Leverington-Waite, Simon Griffiths, Peter R. Shewry, Alison Lovegrove

## Abstract

Increasing dietary fibre (DF) intake is an important target to improve health and an attractive strategy for this is to increase the fibre content of staple foods, particularly white bread which is the staple food in many countries. DF in wheat white flour is derived principally from the endosperm cell wall polysaccharide arabinoxylan (AX) and the water-extractable form of this (WE-AX) accounts for the majority of soluble dietary fibre (SDF), which is believed to confer particular health benefits. We previously identified QTLs for soluble dietary fibre (SDF) on 1B and 6B chromosomes in wheat in biparental populations. Here we show that the 6B high SDF allele encodes a peroxidase protein (PER1-v) with a single missense compared to the more common low SDF form (PER1). Wheat lines with the natural PER1-v allele and with an induced knock-out mutation in PER1 showed similar characteristics of reduced dimerization of ferulate associated with water-extractable WE-AX. Decreased ferulate dimerization is associated with decreased cross-linking of the WE-AX chains and increased solubility of AX. Transiently expressed PER1_RFP fusion driven by native promoter in wheat endosperm was shown to localise to cell walls whereas PER1-v_RFP did not; we therefore propose that PER1-v lacks capacity to dimerise AX ferulate *in vivo* due to mis-localisation. PER1 is the first peroxidase reported to be responsible for oxidative coupling of ferulate on AX, a key process in all grass cell walls. Understanding its role and the effect of variants on AX properties offers a route to control the properties of wheat DF in the human diet.

## Background

Bread wheat is the most widely grown food crop globally and this single species accounts for ~20% of human food calories [www.fao.org/faostat]. Dietary fibre (DF) is an essential component of the human diet and the intake of cereal fibre is associated with a reduced risk of a range of chronic diseases (Gill et al., 2021). The mechanisms of action of fibre are complex and still incompletely understood. However, it has been suggested that the balance of three properties, solubility, viscosity and fermentability in the colon, are important in determining the behaviour of fibre in the GI tract (Gill et al., 2021). Hence, soluble and insoluble forms of fibre share some health benefits but also differ in others.

Wheat is a major source of DF in the western diet, but most products consumed are made from white flour which has a lower content of DF than whole grain (about 4% dry wt. compared with over 10% dry wt.). Increasing the fibre content of white flour is therefore an attractive strategy to deliver health benefits to consumers. The major DF component in white flour is the cell wall polysaccharide arabinoxylan (AX), accounting for about half of the total DF with water-extractable (WE)-AX accounting for most SDF and nearly all of the associated viscosity (Freeman et al., 2016). The amounts of both total AX and WE-AX in white flour vary between wheat genotypes, from about 1.35 to 2.75% dry wt. and 20% and 50% of the total, respectively (Gebruers et al., 2008). Several studies have found QTLs and Marker-Trait Associations (MTAs) for total and WE-AX in wheat (Charmet et al., 2009; Nguyen et al., 2011; Quraishi et al., 2011; Yang et al., 2014; Marcotuli et al., 2015; Yang et al., 2016; Zhan et al., 2019; Lovegrove et al., 2020; Ibba et al., 2021) but no causal genes have been experimentally demonstrated.

Possible candidate genes include those involved in synthesis of AX. AX is a chain of β-1,4-linked xylopyraonsyl units decorated with α-1,3- and α-1,2-linked arabinofuranosyl units; some of α-1,3-linked arabinofuranosyl are themselves 5-O-substituted with ester-linked feruloyl residues. These ferulate moities are key for functionality because they allow for radical oxidative coupling to form ferulate dimers that can cross-link chains of AX (Burr and Fry, 2009; Pellny et al., 2020). This cross-linking is believed to bind AX more tightly into the endosperm cell wall (Saulnier et al., 2007) so decreasing the extent of cross-linking would be expected to increase the amount of WE-AX. Cross-linking of ferulate moities on AX has significance beyond wheat grain as it is a key process in all grass cell walls, determining primary wall extensibility and biomass recalcitrance to digestion and is likely a capability that contributed to evolutionary success of the grasses (Chandrakanth et al., 2023). It has been shown that dimerization of ferulate on AX in grass cell walls requires the action of an apoplastic peroxidase (Burr and Fry, 2009) but the peroxidase gene family is large (Passardi et al., 2004), and the identity of these peroxidases has not been established for any grass cell wall.

Previous studies have identified a QTL for SDF on chromosome 6B in a Yumai34 x Valoris wheat population (Charmet et al., 2009; Lovegrove et al., 2020). Here we show that the causal gene is TraesCS6B02G042500 encoding a peroxidase responsible for cross-linking ferulate moities on AX, decreasing water extractability. The high SDF Valoris allele contains a single missense SNP that appears to prevent localisation of the encoded protein to the cell wall thus disrupting function, resulting in less cross-linking and more WE-AX.

## Results

### RNAseq from endosperm of lines from Yumai34 x Valoris population identifies PER1 as candidate causal gene

We previously mapped significant QTLs for relative viscosity of grain extracts (a property conferred by WE-AX) on chromosomes 1B and 6B and a more minor QTL on 1A in the Yumai34 x Valoris population of doubled-haploid lines (DHLs) (Lovegrove et al., 2020). To identify candidate genes for the 1B and 6B QTLs we analysed RNA-seq from pools of 4 DHLs with contrasting genotypes at these two QTLs (but all with Yumai34 allele at 1A).

Transcripts for enzymes of AX synthesis typically peak around 12-18 days post anthesis (dpa) (Pellny et al., 2012), so we isolated RNA from endosperm at 17 dpa. The 95% confidence intervals [19,946,488 – 28,116,185] for the 6B QTL encompass 323 protein-coding genes annotated in wheat reference genome IWGSC Refseq 1.1; of these 27 were expressed (average FPKM>0.5) and 16 were differentially expressed (FDR<0.05) and/or had splicing differences or coding polymorphisms. From these 16, we identified neighbouring genes TraesCS6B02G042500 and TraesCS6B02G042600 annotated as peroxidases as top candidates due to their expression pattern (high in endosperm during grain fill, not expressed in non-grain tissue) and proximity to the QTL peak. The two genes are result of tandem duplication (sharing 85% CDS identity) and each has 4 exons. RNA-seq mapped to these two genes (Figure 1A) shows that all abundant transcripts are made up of 4 exons but can be from either gene or from TraesCS6B02G042500 exons 1 and 2 combined with TraesCS6B02G042600 exons 3 and 4. Therefore these two separate gene models actually represent splice variants, albeit ones sharing no exons. We named the transcripts and encoded proteins PER1 (identical to TraesCS6B02G042500.1) PER2 (identical to TraesCS6B02G04600.1) and PER1&2 for the hybrid comprised of PER1 exons 1,2 and PER2 exons 3,4. DHLs with the Valoris allele at this 6B locus showed highly significant (P<0.001) differences in expression and splicing with marked decreases in PER1&2 and increases in PER2 expression (Fig. 1B); the Yumai34 sequence is identical to Chinese Spring reference whereas Valoris PER1 has 3 synonymous SNPs and one missense SNP in exon 1 of PER1. The missense results in a Ser51Phe change in the protein encoded by the Valoris allele which we call PER1-v. Both PER1 and PER2 have homeologues on the A and D subgenomes but PER1A is not expressed, and the other forms are less expressed than PER1 and PER2 (Fig. 1B). We hypothesised that, despite the likely redundancy between these forms, the missense and/or lower expression of PER1-v in Valoris might be the cause of the higher SDF.

**Figure 1.**
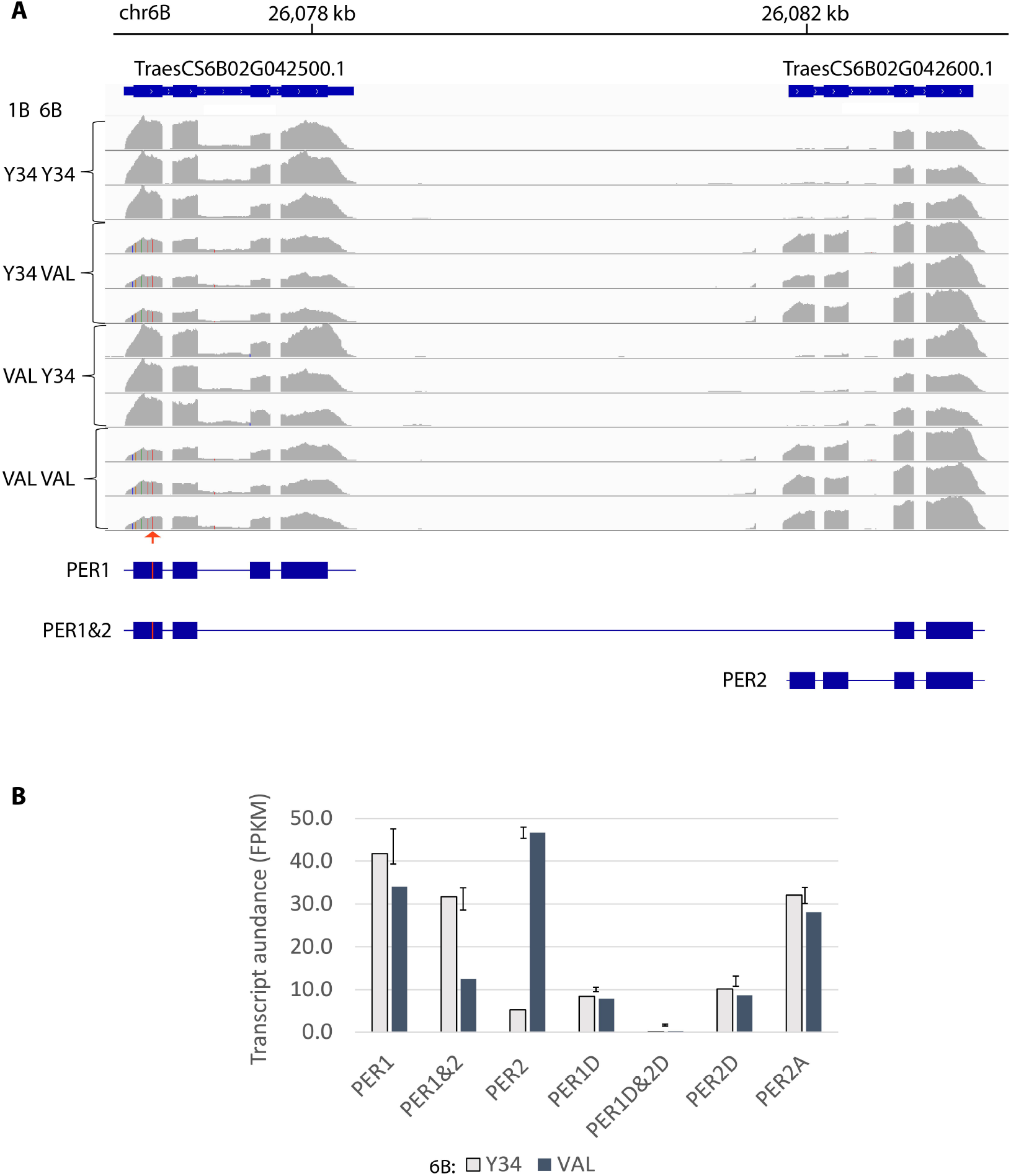
RNAseq analysis of PER transcripts from 17 dpa endosperm of DHLs in Yumai34 x Valoris mapping population. A. Read coverage for 6B chromosome region containing TraesCS6B02G042500 and TraesCS6B02G042600 genes from 12 samples of DHLs selected to have Yumai34 (Y34) or Valoris alleles (VAL) at 1B and 6B QTLs with 3 reps of each combination. The arrow indicates the missense SNP in Valoris. Analysis of paired reads showed nearly all transcripts were one of the 3 forms depicted PER1, PER1&2 or PER2. These inferred transcripts are shown as blocks and lines where blocks represent coding exons and the missense SNP in Valoris allele is marked in red. B. Transcript abundance of the splice variants PER1, PER1&2 or PER2 and their homeologues on sub-genomes A and D. PER1A exons are not expressed. Bar heights and error bars are estimated means and 5% LSD for 6B alleles of each transcript from a 2-way ANOVA of 1B and 6B allele effects. No 1B effects were significant (P>0.05) so results shown group DHLs differing in 1B alleles (n=6).

### DHLs with Valoris allele at 6B have lower dimerization of ferulate in endosperm WE-AX

The identification of PER1 as candidate causal gene suggested that the mode of action for the 6B QTL could be via the dimerization of ferulate on AX, with lower dimerization associated with the Valoris PER1-v allele. We have previously determined dimerization of wheat endosperm AX ferulate in total and WE endosperm fractions and increases were seen in both when genes responsible for synthesis of the AX xylan backbone were downregulated (Freeman et al., 2017; Pellny et al., 2020). Here, we observed significantly lower ferulate dimerization on WE-AX in DHLs with the Valoris genotype (6B:H) to those with the Yumai34 genotype (6B:L) in the 6B QTL region (Table 1). Out of the five ferulate dimer (diFA) peaks detected, the ratios to FA monomer were all decreased in 6B:H DHLs and the sum of all dimers expressed as % dimerization was significantly decreased (F. prob = 0.014) by ~30%. The effects of 1B:H were more complex, decreasing some dimers significantly but having no significant effect on overall dimerization. There were also significant effects of 1B and 6B genotypes and interaction of these on FA monomer amount per unit dwt endosperm (Table 1).

**Table 1.** Amounts of ferulate monomer (FA) and dimers (diFA) in WE fractions from endosperm of DHLs grouped according to genotype at 1B and 6B QTLs. Results from a 2-way ANOVA of 1B x 6B genotype showing means for each group (n=3) and F probability for effects.

|  |  | diFA / FA (w/w) |  |  |  |  |  |  |
| --- | --- | --- | --- | --- | --- | --- | --- | --- |
| means |  | Dimerisation<br>[tot diFA /<br>(tot diFA +<br>FA)] | diF8-8AT | diF8-8 | diF8-5 | diF5-5 | diF8-0-4<br>& diF8-5BF | FA (µg / g<br>dwt) |
|  | 1B:L 6B:L | 22% | 0.065 | 0.024 | 0.046 | 0.08 | 0.061 | 6.3 |
|  | 1B:H 6B:L | 19% | 0.038 | 0.072 | 0.032 | 0.048 | 0.049 | 12.9 |
|  | 1B:L 6B:H | 15% | 0.032 | 0.017 | 0.023 | 0.058 | 0.05 | 10.5 |
|  | 1B:H 6B:H | 14% | 0.032 | 0.015 | 0.027 | 0.046 | 0.04 | 13.4 |
| F prob. | 1B | 0.319 | 0.11 | 0.405 | 0.665 | 0.004 | 0.024 | <.001 |
|  | 6B | 0.014 | 0.026 | 0.251 | 0.266 | 0.072 | 0.041 | 0.013 |
|  | interaction | 0.794 | 0.09 | 0.362 | 0.477 | 0.125 | 0.755 | 0.035 |

**Table 2.** Amounts of ferulate monomer (FA) and dimers (diFA) in endosperm of Cadenza BC2F2 lines segregating for KO mutation in PER1. Results from a 1-way ANOVA of mutation showing means for each group (n=7) and F probability for effect.

|  |  | diFA / FA (w/w) |  |  |  |  |  |  |  |
| --- | --- | --- | --- | --- | --- | --- | --- | --- | --- |
| means |  | Dimer-<br>isation<br>[tot diFA /<br>(tot diFA<br>+ FA)] | diF8-8AT | diF8-8 | diF8-5 | diF5-5 | diF8-0-4 | diF8-5BF | FA (µg<br>/ g<br>dwt) |
|  | mutant | 15.5% | 0.018 | 0.036 | 0.049 | 0.027 | 0.052 | 0.053 | 88.8 |
|  | null | 17.0% | 0.024 | 0.04 | 0.05 | 0.035 | 0.056 | 0.061 | 79.3 |
| F prob. | mutation | 0.037 | 0.019 | 0.227 | 0.830 | 0.030 | 0.346 | 0.046 | 0.194 |

### Wheat lines with an induced mutation in PER1 have lower dimerization of ferulate in endosperm

Using the mutagenised, exome-sequenced population of wheat var. Cadenza (Krasileva et al., 2017), we identified a line carrying a premature stop codon in exon 2 of PER1. We twice backcrossed lines to Cadenza wild-type and allowed lines to self-fertilise for 2 further generations (BC2F2 lines) and identified segregants that were homozygous for the mutation and those that were homozygous for the wild-type allele. We then compared the ferulate dimerization in endosperm tissues from these; BC2F2 lines carrying the mutation had significantly (F prob. = 0.037) lower ferulate dimerization in endosperm (Table 2) supporting the putative role of PER1.

### PER1_RFP fusion protein transiently expressed in wheat endosperm localises partially to cell wall but PER1-v_RFP does not

To investigate whether the missense SNP could explain the effect of PER1-v allele (as opposed to the expression difference; Fig. 1B) we made constructs with and without the missense SNP to encode PER1 and PER1-v, both fused at C-terminal to RFP reporter and driven by the endogenous promoter. We transiently transformed wheat endosperm tissue with plasmids containing these constructs. For both constructs, the RFP signal was mostly in the cytoplasm, possibly as a result of high expression overloading the capacity of the cell to normally process the protein which is predicted to require signal peptide cleavage and N-glycosylation. However in plasmolysed cells expressing PER1_RFP we also observed RFP signal associated with the cell wall whereas in plasmolysed cells expressing PER1-v_RFP we never observed cell wall RFP signal (examples shown in Figure 2, the complete set of images with clearly plasmolysed cells are in Supplemental Figures S1-S4). In total, we observed cell wall RFP signal in 41 out of 43 plasmolysed cells with PER1_RFP whereas there were none in 40 plasmolysed cells expressing PER1-v_RFP). Since we would expect most or all AX ferulate dimerization to occur in the cell wall (Burr and Fry, 2009), the failure of PER1-v to localise to the cell wall means it is likely defective in function.

**Figure 2.**
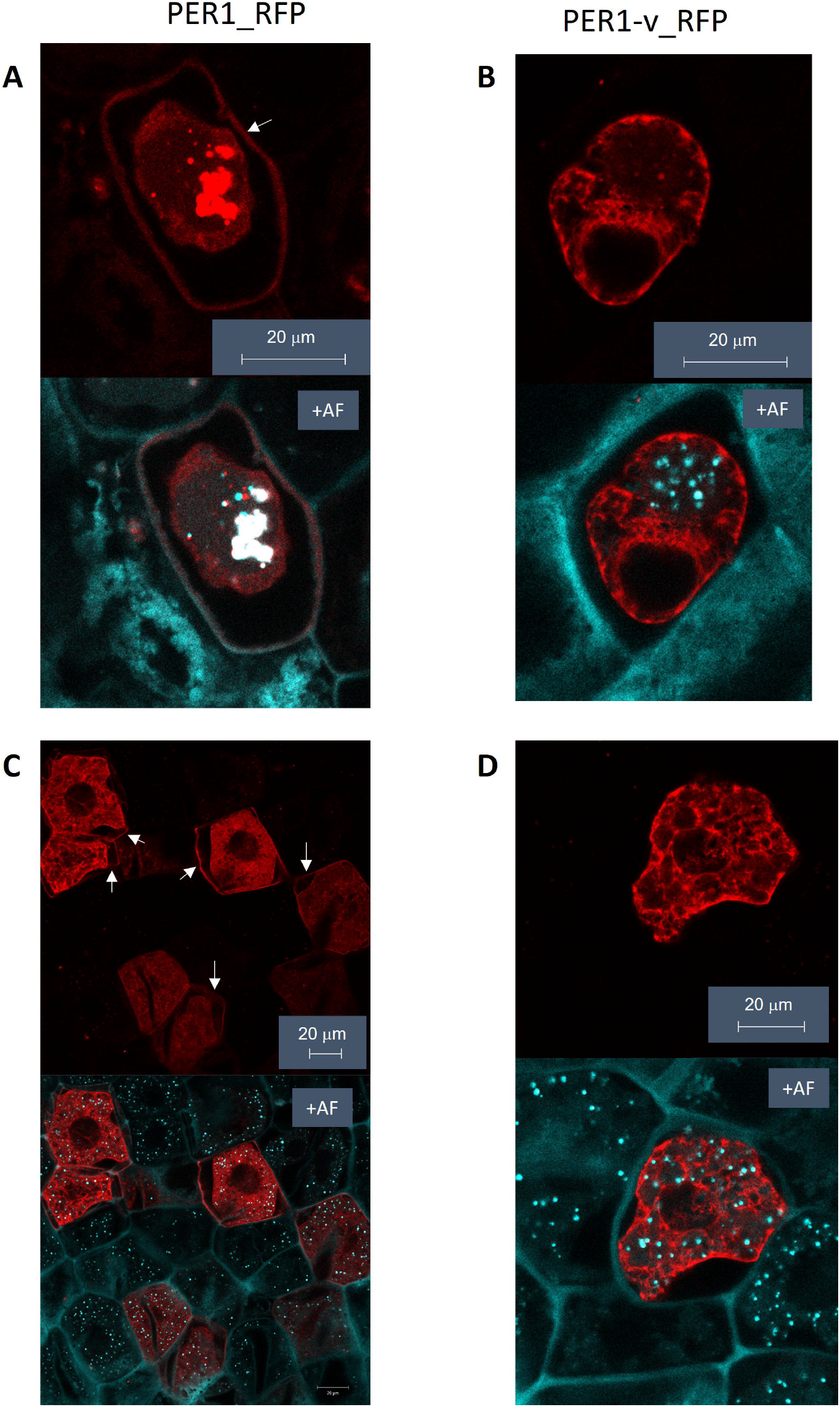
Example images of sections of plasmolysed wheat endosperm cells transiently expressing PER1_RFP and PER1-v_RFP proteins. Pairs of images show RFP fluorescence signal alone in top panel and combined with autofluorescence (+AF) in lower panel to show cell walls. Plasmolysis caused complete **(A, B)** or partial **(C,D)** separation of cell contents from cell wall allowing identification of RFP signal from cell wall indicated by arrows. Cell wall localisation of RFP signal is present in cells expressing PER1_RFP **(A, C)** but not those expressing PER1-v_RFP **(B, D).**

### Occurrence of missense SNP in wheat varieties

The effect of the single missense SNP on PER1 localisation strongly suggests that this is the cause of the 6B QTL for SDF. If this is the case, this SNP which is chr6B 26076727 C->T on IWGSC 1.0 reference (Axiom marker AX-94816599) represents a perfect marker for the trait. This SNP was also one of 5 closely neighbouring SNPs on 6B found by GWAS as associated with equal significance to high relative viscosity of aqueous extracts in the WHEALBI panel (Lovegrove et al., 2020). The increasing allele found in Valoris is not rare, being present in 150 out of 426 panel members (www.whealbi.eu). We further investigated allele frequency in another panel (an elite high fibre panel, comprised of advanced commercial breeding lines of UK, French and German winter wheats) using a KASP marker designed to the SNP. In this panel, we found the Valoris allele in 57 lines, 287 giving reference allele and 27 scored as het.

## Discussion

Soluble dietary fibre may confer health benefits via several mechanisms including fermentation to release short-chain fatty acids in the colon, and by increasing viscosity which may reduce the rate of glucose release from starch in the duodenum (Gill et al., 2021). In the case of wheat SDF, nearly all of the viscosity of aqueous extracts of white flour is due to WE-AX (Freeman et al., 2016). AX is a major component of all grass cell walls, but is normally not extractable in water, so wheat endosperm WE-AX is unusual in this respect. Extractability of AX in water is determined by several structural factors of which cross-linking via ferulate dimerization may be the most important (Saulnier et al., 2007). The AX in the starchy endosperm, and particularly the WE-AX fraction, has less ferulate [AX ~0.5%; WE-AX ~0.2% (Freeman et al., 2017)], and therefore potential for cross-linking, than AX from other tissues in wheat and in grasses generally, which varies greatly but can be up to 20% (Hatfield et al., 2017).

It has been shown that oxidative coupling of ferulate that results in dimers and higher oligomers bound to AX in grass cell walls occurs in the apoplast and is mediated by peroxidases (Burr and Fry, 2009) but these have not been identified. Here we provide strong evidence that the dimerization of ferulate on wheat endosperm AX is mediated by the PER1 protein, since both a natural allele and an induced KO mutant result in lower dimerization (Tables 1 and 2). The natural allele has a missense SNP that prevents localisation of PER1 to the cell wall (Fig. 2), the likely site for AX dimerization. We therefore attribute the 6B QTL effect on SDF to this SNP resulting in a defective PER1 that lowers the capacity for AX ferulate dimerization, resulting in a higher proportion of AX being extractable in water. This conclusion is clear despite the changes in expression of PER1, PER1&2 and PER2 associated with the Valoris 6B genotype (Fig. 1) which could be due to nearby *cis* factors rather the missense SNP itself.

PER1 is a member of a large gene family of class III haem peroxidases that are specific to green plants. These peroxidases apparently all catalyse the same reaction *in vitro,* generating radical oxygen species from H_2_O_2_ and it has been hypothesised that *in vivo* specificity is achieved by localisation close to the intended substrate (Francoz et al., 2015). This is supported by observation of precise localisation of peroxidases associated with lignification in Arabidopsis, (Hoffmann et al., 2020) and has been demonstrated for a peroxidase that binds to a particular cell wall microdomain on the polysaccharide homogalacturonan in order to function in mucilage extrusion (Francoz et al., 2019) and for a lignin peroxidase that localises to the Casparian strip via binding to another protein (Lee et al., 2013). It is therefore probable that a large family of peroxidase genes is required for the multiplicity of peroxidase binding sites that confer specificity of function. This is consistent with our result showing that PER1 is present in the cell wall whilst PER1-v is not and this difference is associated with decreased dimerization of AX ferulate. The single amino acid difference between the natural and mutant peroxidases is at a serine residue in PER1 which is predicted by AlphaFold to be involved in H-bonding in an α-helix on the exterior of the protein (Figure S5). This is changed in PER1-v to the aromatic amino acid phenylalanine which would prevent H-bonding and disrupt the α-helix. PER1 and PER2 are in a grass-specific subclade of Group VI of the class III peroxidase gene family as defined in (Passardi et al., 2004) (Figure S6). This is consistent with a specific role in cross-linking of AX ferulate, which is confined to grasses and other commelinid monocots (Chandrakanth et al., 2023). PER1 and PER2 are only expressed in grain out of the major wheat tissues (Figure S5) suggesting they may have specific properties related to the unusual structure of grain AX.

Our identification of PER1 as the cause of the high SDF trait from the 6B QTL will allow the specific tailoring of the solubility of wheat grain AX to increase benefits for human health. It also represents an important step forward in our broader understanding of grass cell walls.

## Methods

### Plant Growth

We grew the Yumai34 x Valoris population in the field in 2018 and in the glasshouse in 2022 in randomised designs at Rothamsted Research, UK. In both experiments, we selected DHLs as having the same genotype for all Axiom genotyping markers on 35K SNP platform within the 1A, 1B, 6B QTLs and assigned these to 4 pools of contrasting parental genotype at 1B and 6B but all with Yumai34 genotype at 1A. Three biological reps of these pools from different blocks were used from 2018 field experiment for transcriptome analysis and from 2022 glasshouse experiment for ferulate determination.

### KASP marker for Valoris missense SNP

We designed KASP marker to Valoris chr6B 26076727 SNP and used this to confirm SNP genotype of the selected lines in Yumai34 x Valoris population and in elite high fibre panel. Primers and PCR conditions are given in Table S1.

### RNA-seq analysis

Total RNA was isolated from developing endosperm of 20 selected DHLs at 17 days post anthesis as previously described (Pellny et al., 2012; Pellny et al., 2020) to give 3 biological replicates of each pool. Library preparation and mRNA sequencing was done by Novogene (HK, China). We mapped reads to IWGSC refseq 1.1 genome using HISAT2 (Kim et al., 2015), visualised alignments using Integrative Genomics Viewer (Robinson et al., 2011) and estimated transcript abundance and differentially expressed genes using DESeq2 (Love et al., 2014). Transcript abundances for TraesCS6B02G042500.1 and TraesCS6B02G042600.1 gene models in reference were reallocated to PER1, PER1&2 and PER2 based on manually determined ratios of counts of read pairs that could be clearly assigned to these.

### Ferulate Monomer and Dimer Analysis

We analysed the ferulate monomer and dimer content of endosperm fraction from mature grain by HPLC analysis as previously described (Freeman et al., 2017) using pure ferulate dimer standards kindly supplied by Professor John Ralph (Lu et al., 2012).

### PER1 Knock-Out Mutant

From the database of mutations in the exome sequenced collection of mutagenised wheat cv Cadenza available at Ensembl Plants, we identified line Cadenza1644 carrying a G->A mutation at 26,076,941 on chr6B causing a premature stop codon Trp94STOP in exon 2 of PER1. We backcrossed this line twice to Cadenza, allowed to self-fertilise over two further generations and identified BC2F2 homozygotes for both mutation and wild-type allele using crossing and genotyping procedures previously described (Pellny et al., 2020); subgenome specific primers for genotyping are given in Table S1. We grew these lines in a randomised design in the glasshouse with 7 replicate pots per line.

### Transient expression of PER1_RFP and PER1-v_RFP in immature wheat endosperm

We designed a version of PER1 CDS (TraesCS6B02G042500.1) that contained additional cloning sites immediately before the natural NcoI site present over the ATG codon and an HpaI site at the C’ terminal end of the gene that involved the addition of an extra serine residue to the protein. Nine synonymous changes were also engineered into the coding sequence to facilitate cloning while retaining the natural PER1 protein sequence. We designed PER1-v from this with a single bp change corresponding to the Valoris allele missense SNP giving rise to the Ser51Phe peptide sequence change. From alignment analysis of the 5’ upstream sequences of PER1 and the Chinese Spring, A, and D homeologues and other wheat relatives, we selected 2135 bp of the PER1 sequence as likely to contain a complete promoter including the 5’ transcribed untranslated sequences. Bases immediately upstream of the ATG codon were modified to contain the same restriction sites as engineered into the PER1 and PER1-v genes sequences for cloning purposes and to remove the out of frame ATG codon present 5bp upstream of the PER1 start codon. The designed PER1, PER1-v and PER1 promoter sequences were synthesised by Genscript (Oxford, UK) and transferred using standard cloning techniques to pRRes208, a biolistic vector backbone. Two plasmids were generated with the PER1 promoter driving either PER1 [pRRes208.618] or PER1-v [pRRes208.617]. In each construct the PER1 sequences were fused in frame upstream of a six amino acid linker sequence and the fluorescence marker gene Tag-RFP_T. Immature wheat endosperm sections were isolated from 7-10 dpa wheat seeds cv Cadenza, as described in (Jones et al., 2008). The tissues were bombarded on the same day as isolation using 0.6μm gold particles (Bio-Rad Laboratories Ltd, UK) coated with the constructs pRRes208.617, pRRes208.618 or pRRes.380, a plasmid containing Tag-RFP under the control of the constitutive rice actin promoter which acted as a positive control. Bombardment was carried out using the PDS-1000/He particle gun with 650psi rupture pressure and 29” Hg vacuum; full details of bombardment parameters can be found in (Sparks and Doherty, 2020). Plates were incubated in the dark at 22°C for 1 day prior to bioimaging.

### Imaging of wheat endosperm transiently expressing PER1_RFP and PER1-v_RFP

Transformed, immature wheat endosperm sections were mounted on glass slides with coverslips for imaging with a Zeiss LSM 780 confocal microscope (Carl Zeiss Ltd. Cambourne, Cambridge, UK). The endosperm tissue was mounted in distilled water for initial imaging and the water replaced under the coverslip with 2M NaCl by capillary action to induce plasmolysis. Cells were monitored for up to 10 mins to observe plasmolysis and imaged with the 40x objective using 405nm excitation, 428nm – 490nm emission for cell wall auto-fluorescence, and 561nm excitation, 580nm – 700nm emission for Tag-RFP.

## Supplementary material

Figures S1. Images from endosperm grain sections transiently expressing PER1_RFP and PER1-v_RFP where clear cell plasmolysis was observed. Pairs of images show RFP fluorescence signal alone in top panel and combined with autofluorescence (+AF) in lower panel to show cell walls. Plasmolysis separation of cell contents from cell wall allowing identification of RFP signal from cell wall indicated by arrows. Cell wall localisation of RFP signal is present in cells expressing PER1_RFP **(A, B, E, F, I, J)** but not those expressing PER1-v_RFP **(C, D, G, H, K, L**).

Figures S2. Further images from endosperm grain sections transiently expressing PER1_RFP and PER1-v_RFP where clear cell plasmolysis was observed. Details as for Fig. S1, except for image pairs **(D, G, H, K, L)** where image in upper panel is pre-plasmolysis and after plasmolysis in lower panel.

Figures S3. Further images from endosperm grain sections transiently expressing PER1_RFP and PER1-v_RFP where clear cell plasmolysis was observed. Details as for Fig. S1, except for image pairs **(C-L)** where image in upper panel is pre-plasmolysis and after plasmolysis in lower panel. Image pair **F** is the single exception where we could not see clear cell wall localisation of RFP signal in cells expressing PER1_RFP.

Figures S4. Further images from endosperm grain sections transiently expressing PER1_RFP and PER1-v_RFP where clear cell plasmolysis was observed. Pairs of images show pre-plasmolysis image in upper panel and after plasmolysis in lower panel. Plasmolysis causes separation of cell contents from cell wall allowing identification of RFP signal from cell wall indicated by arrows. Cell wall localisation of RFP signal is present in cells expressing PER1_RFP **(A)** but not those expressing PER1-v_RFP (**B, C, D).**

Figure S5. AlphaFold predicted structure of PER1 (TraesCS6B02G04600.1) protein with Ser51 residue that is changed in PER1-v highlighted. Predicted N-glycosylation Asn, haem-binding and signal peptide cleavage sites are indicated.

Figure S6. Phylogeny of subclade within class III peroxidase family containing PER1 and PER2 genes. For phylogenetic analyses of protein sequences, we selected the rice ortholog of PER genes (OsPRX91) and its three closest paralogs (OsPrx90, OsPrx92, OsPrx93) which are all neighbouring genes on rice chromosome 6. We then selected all orthologs of these from wheat, Arabidopsis, *Ananas comosus, Glycine max, Hordeum vulgare, Populus trichocarpa* and *Sorghum bicolor* as defined Ensembl Plants release 56 (Yates et al., 2021) and aligned with MUSCLE (Edgar, 2004) and generated phylogenetic tree with Phyml (Guindon et al., 2010) as previously described (Pellny et al., 2012). Wheat expression data was taken from wheat-expression.com (Borrill et al., 2016) averaged across 34 studies encompassing many genotypes and environments.

