## Supplemental Figures S1-S4 for "A high soluble-fibre allele in wheat encodes a defective cell wall peroxidase responsible for dimerization of ferulate moieties on arabinoxylan"

### Slide 1
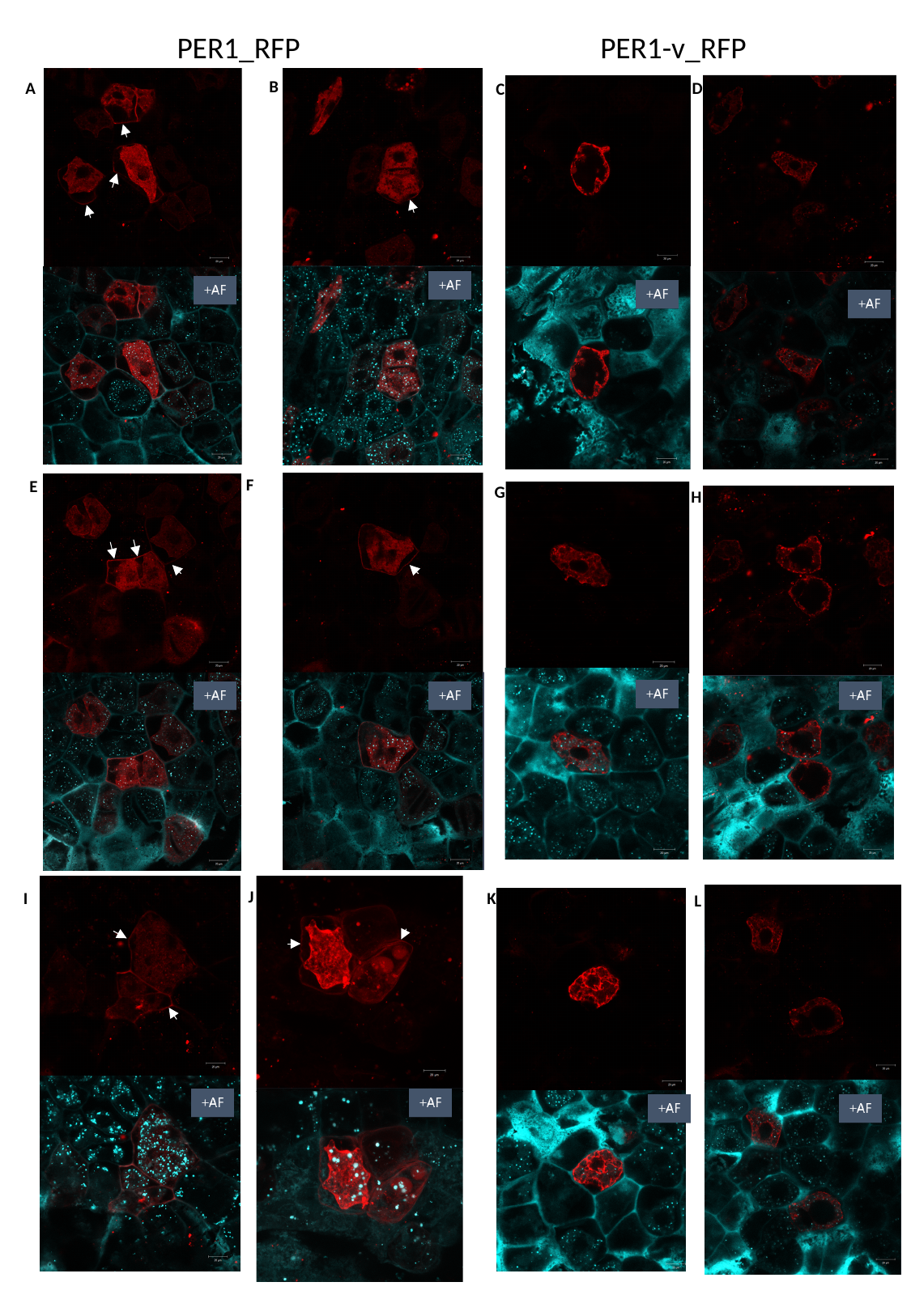

PER1_RFP
PER1-v_RFP
B
A
D
C
F
E
G
H
J
I
K
L

### Slide 2
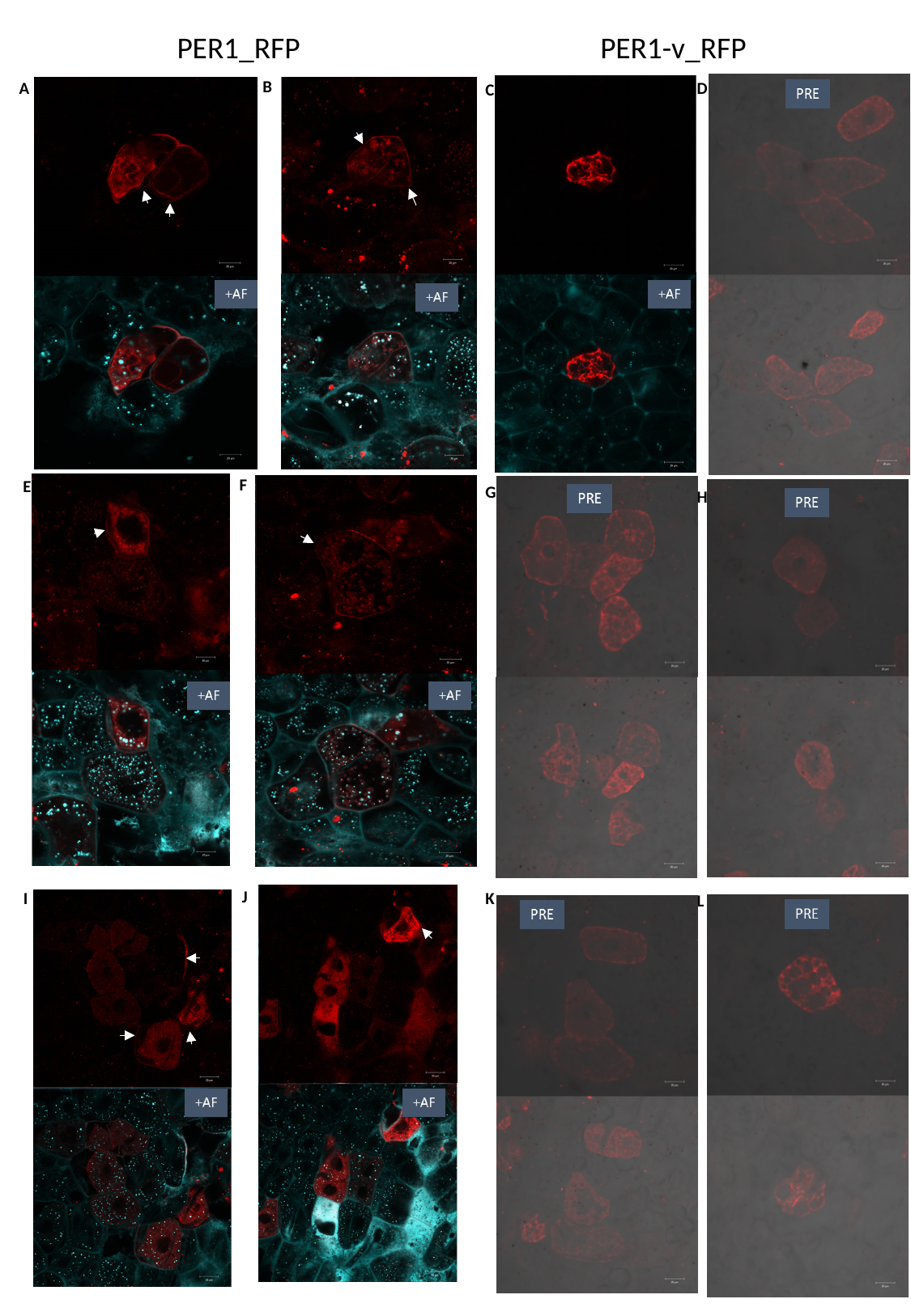

PER1_RFP
PER1-v_RFP
B
A
D
C
F
E
G
H
J
I
K
L

### Slide 3
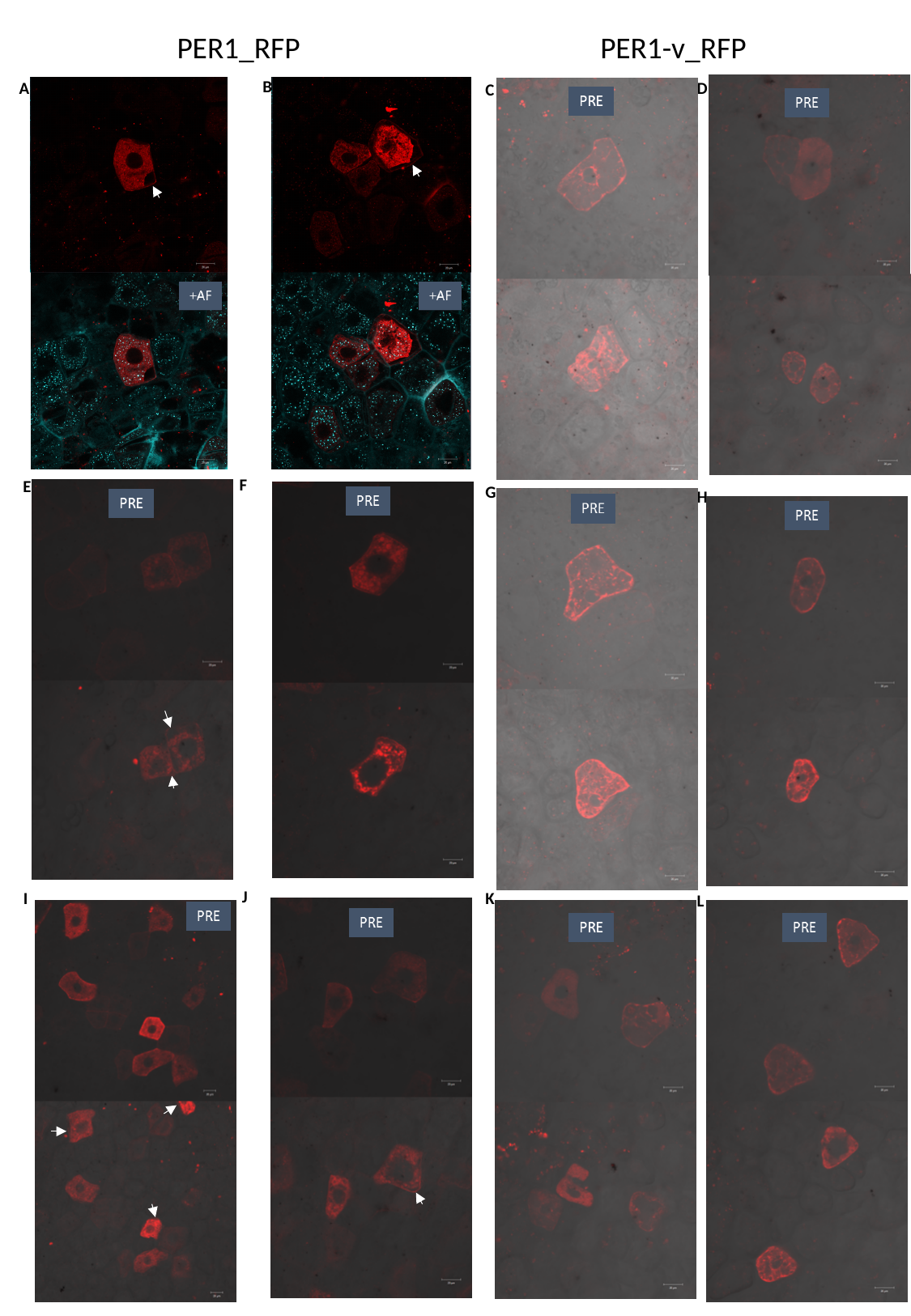

PER1_RFP
PER1-v_RFP
B
A
D
C
F
E
G
H
J
I
K
L

### Slide 4
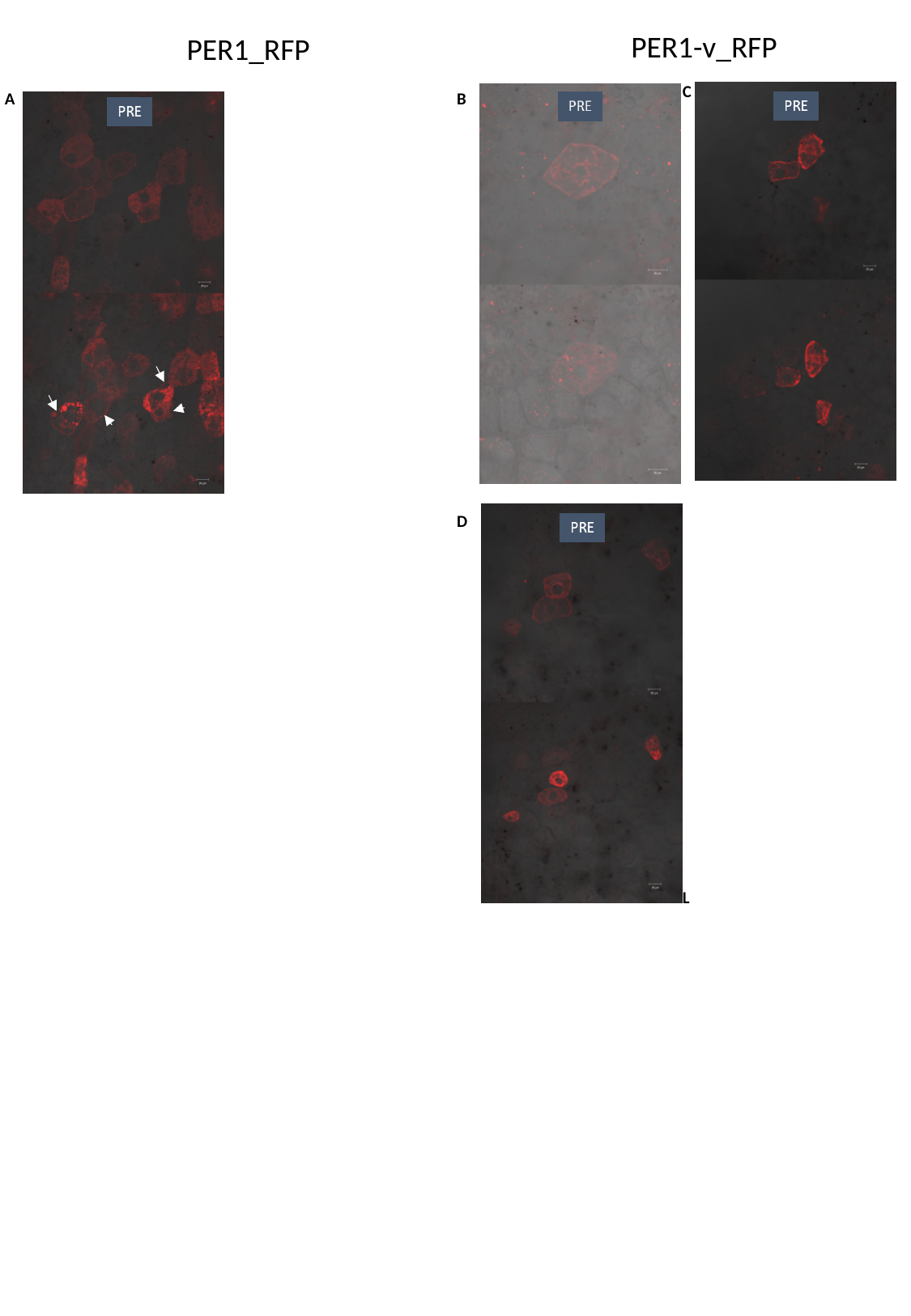

PER1-v_RFP
PER1_RFP
C
A
B
D
L
