## Supplementary figures and images for "A high soluble-fibre allele in wheat encodes a defective cell wall peroxidase responsible for dimerization of ferulate moieties on arabinoxylan"

### Supplemental Figure S5

#### Slide 1
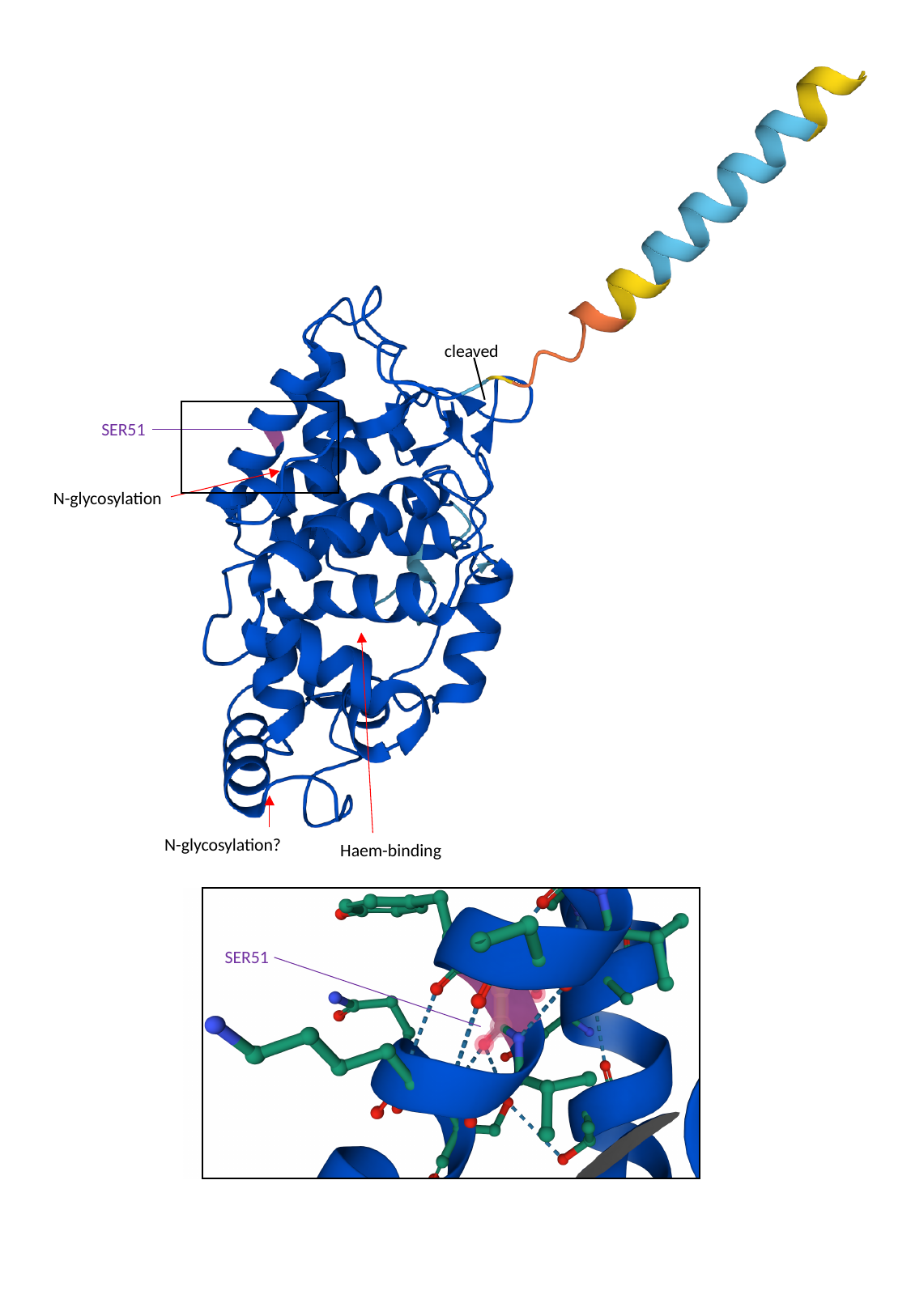

cleaved
SER51
N-glycosylation
N-glycosylation?
Haem-binding
SER51

### Supplemental Figure S6

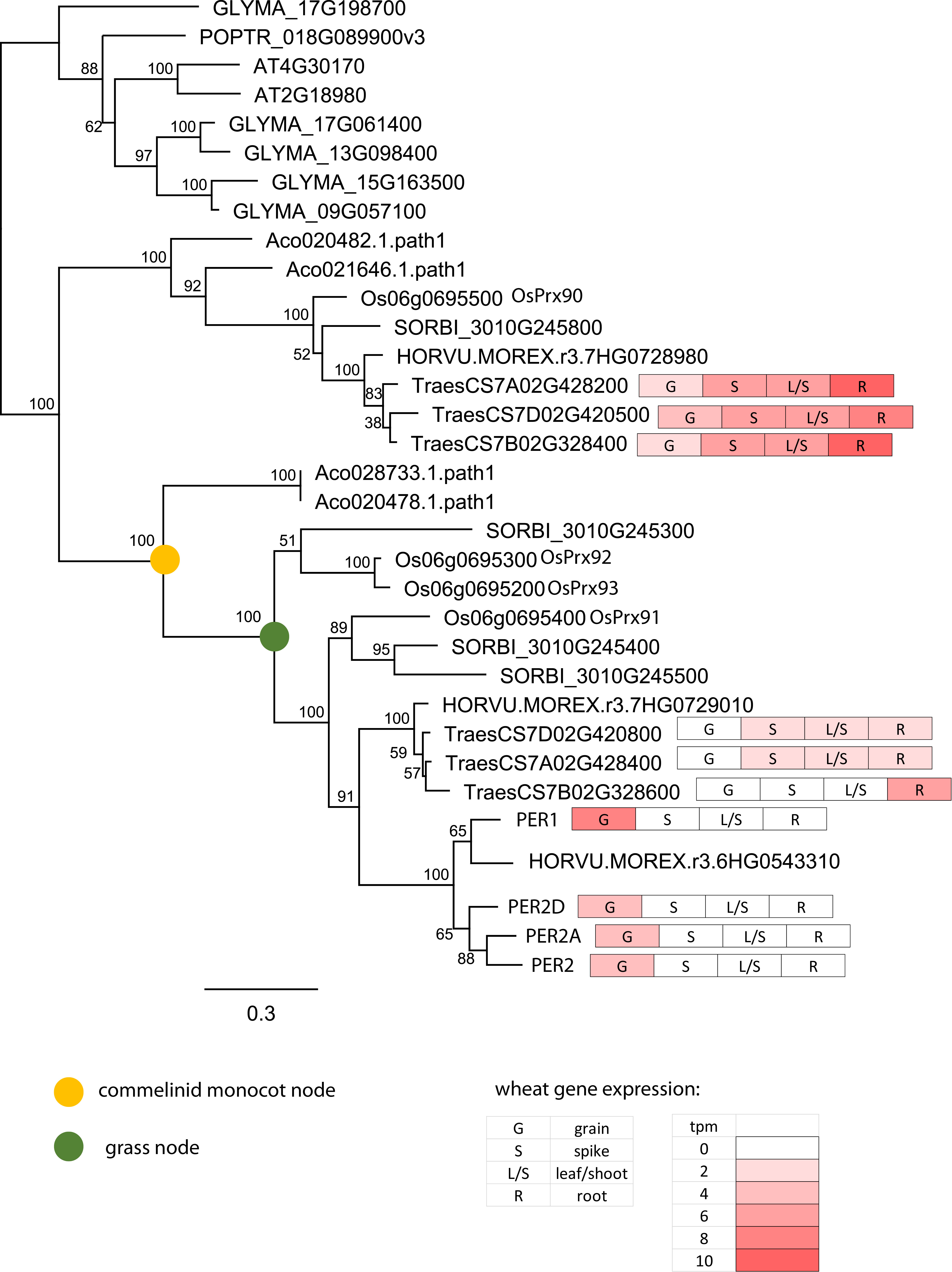
